# Improving the flexibility of Genetically Encoded Voltage Indicators via intermolecular FRET

**DOI:** 10.1101/2020.07.31.230128

**Authors:** Lee Min Leong, Bok Eum Kang, Bradley Baker

## Abstract

A new family of Genetically Encoded Voltage Indicators (GEVIs) has been developed based on inter-molecular Förster Resonance Energy Transfer (FRET). To test the hypothesis that the GEVI, ArcLight, functions via interactions between the fluorescent protein (FP) domain of neighboring probes, the FP of ArcLight was replaced with either a FRET donor or acceptor FP. We discovered relatively large FRET signals only when cells were co-transfected with both the FRET donor and acceptor GEVIs. Using a CFP donor and an RFP acceptor, we were able to observe a voltage dependent signal with a Stokes shift of over 200 nm. The intermolecular FRET strategy also works for rhodopsin-based probes potentially improving their flexibility as well. Separating the FRET pair into two distinct proteins has important advantages over intramolecular FRET constructs. First, the signals are larger. Apparently the voltage-induced conformational change moves the two FPs independently thereby increasing the dynamic range. Second, the expression of the FRET donor and acceptor can be restricted independently enabling greater cell type specificity as well as refined subcellular voltage reporting.

## Introduction

The ability of Genetically Encoded Voltage Indicators (GEVIs) to optically report changes in membrane potential offers the promise of monitoring neuronal activities from multiple cell populations in neuronal circuits simultaneously. Despite the impressive progress in the development of GEVIs (Adam et al., 2019; Chamberland et al., 2017; Kannan et al., 2018; Lee et al., 2017; Marshall et al., 2016; Piatkevich et al., 2018; Yi, Kang, Lee, Braubach, & Baker, 2018), this multiple observation potential remains largely theoretical. A recent, independent comparison of GEVIs revealed that while most perform well in cultured, single cell recordings, the only consistent signal observed *in vivo* was from ArcLight (Bando, Sakamoto, Kim, Ayzenshtat, & Yuste, 2019). ArcLight’s brightness, dynamic range, and voltage sensitivity makes it one of the easiest GEVIs to use. However, the need to average trials indicates that ArcLight’s signal also needs improvement.

In order to enhance the optical signal of ArcLight, we have sought to elucidate the mechanism mediating the voltage-dependent fluorescence change upon plasma membrane potential fluctuations. ArcLight consists of a pH sensitive fluorescent protein (FP), Super Ecliptic pHluorin A227D (SEpH), (Ng et al., 2002) fused to the voltage sensing domain (VSD) of the *Ciona intestinalis* voltage sensing phosphatase (Jin et al., 2012). The SEpH FP in ArcLight contains a unique mutation that introduces a negative charge to the exterior of the β-can structure. A previous report from our lab suggested that this external negative charge interacts with a neighbouring FP since mutations to SEpH that favour the monomeric form of the FP diminished the voltage-dependent signal by at least 70% (Kang & Baker, 2016). Taking advantage of this FP dimerization architecture, we were able to develop the red-shifted GEVI, Ilmol, by replacing SEpH with the FP, dTomato (Yi et al., 2018).

To test the hypothesis that the FP domains of neighboring probes for ArcLight-type GEVIs are capable of interacting, we replaced SEpH in ArcLight with FPs capable of Fluorescence Resonance Energy Transfer (FRET). Previous FRET versions of GEVIs have used intramolecular FRET where the donor and acceptor chromophore are contained in the same protein (Akemann et al., 2012; Dimitrov et al., 2007; Lundby, Akemann, & Knopfel, 2010; Sung et al., 2015). Here we separate the FRET pair into two distinct proteins by fusing the donor FP to one voltage-sensing domain (VSD) while the acceptor is fused to a separate VSD. Co-transfection of ArcLight-derivatives when SEpH was replaced by the cyan fluorescent protein (CFP), Cerulean, or the yellow fluorescent protein (YFP), Venus, resulted in robust FRET signals. These FRET signals prove that the FP domains of ArcLight are capable of interacting. Furthermore, interFRET-GEVIs consisting of green/red FRET pairs also yielded robust signals indicating that the entire visible spectrum is open for future probe development as well as the potential for far-red/infra-red FRET pairs.

This new family of Inter-FRET GEVIs offers several advantages over previously reported probes. The ratiometric nature of the FRET signal enables the removal of motion artefacts due to blood flow and respiration during *in vivo* recordings. Inter-FRET GEVIs will also improve 2-photon probe development since any bright 2-photon FP can now be used as the donor in conjunction with an appropriate FRET acceptor FP. Cell type specificity may also be improved since different promoters can be used to express the donor and acceptor partners. Only cell types that express both promoters will be capable of yielding a voltage-dependent FRET signal. As a proof of principal, we were also able to elicit a voltage-dependent optical signal from a rhodopsin-based GEVI via intermolecular FRET.

## Materials and Methods

### Plasmid design and construction

ArcLight-Cerulean, ArcLight-Venus, ArcLight-Clover, ArcLight-mRuby2, Bongwoori-R3-Cerulean, Bongwoori-R3-Venus, Bongwoori-R3-dTomato, CC1-Cerulean, and CC1-Venus were constructed by replacing the SE227D FP in ArcLight with the respective FPs. The farnesylated versions of the FPs were synthesised (Integrated DNA Technologies, USA). For the ArcLight and Bongwoori-R3 constructs, the FPs were designed to have a 5’ BamHI restriction site and a 3’ stop codon followed by a XhoI site. The FPs for the CC1 constructs were designed to have a 5’ BamHI restriction site and a 3’ stop codon followed by a XhoI site. The Ace2N construct was designed to have a 5’ NheI site and a 3’ stop codon followed by a XhoI site. Conventional one-step and two-step PCR overlap PCR were used to generate monomeric versions of FPs and the Ace2N construct. Primers were designed to introduce the restriction sites and the monomeric mutations in the FPs. The backbone vector used in this study is pcDNA3.1 with a CMV promoter.

### Cell culture and transfection

HEK 293 cells were cultured in Dulbecco’s Modified Eagle Medium (DMEM; Gibco, USA) supplemented with 10% Fetal Bovine Serum (FBS; Gibco, USA) in a 37 °C, 100% humidity and 5% CO_2_. For transfection, HEK 293 cells were suspended using 0.25% Trypsin-EDTA (Gibco, USA) then plated onto poly-L-lysine (Sigma-Aldrich, USA) coated #0 coverslips (0.08-0.13mm thick and 10 mm diameter; Ted Pella, USA) at 70% confluency. Transient transfection was carried out with Lipofectamine 2000 (Invitrogen, USA) according to the manufacturer’s protocol. Equal concentrations of DNA were used for co- and tri-transfection experiments.

Hippocampal neurons were cultured according to an approved animal experiment protocol by the Institutional Animal Care and Use Committee at KIST (animal protocol 2016-082) as described previously (Lee et al. 2017). In brief, the hippocampi of embryonic day 17 C57BL/6 mice (Koatech, South Korea) were dissected and digested with 0.125% Trypsin-EDTA solution (Gibco, USA) for 15 min in a 37 °C water bath. DNase I (Sigma Aldrich, USA) was added for 30 sec after digestion. The cells were then rinsed with plating media (PM) composed of 10% FBS and 1% penicillin-streptomycin (Gibco, USA) in DMEM and subjected to mechanical trituration. Dissociated neurons were then plated onto poly-D-lysine (Sigma-Aldrich, USA) coated #0 coverslips at 5 × 10^4^ cells/ml density. The PM was replaced with maintenance media (MM) composed of Neurobasal medium (Gibco, USA), supplemented with 2% B27 supplement (Gibco, USA) and 1% penicillin-streptomycin. 50% of MM was exchanged every 3 days. Transient transfection of cultured mouse hippocampal neurons were done 5-7 days in vitro (DIV) using Lipofectamine 2000 according to the manufacturer’s protocol and experimented on DIV 8–12. Equal concentrations of DNA plasmids were used for co-transfection experiments.

### Electrophysiology

Coverslips with transiently transfected cells were inserted into a patch chamber (Warner instruments, USA) with its bottom side covered with a #0 thickness cover glass for simultaneous voltage clamp and fluorescence imaging. The chamber was kept at 34 °C throughout the experiment and perfused with bath solution (150 mM NaCl, 4 mM KCl, 1 mM MgCl_2_, 2 mM CaCl_2_, 5 mM D-glucose and 5 mM HEPES, pH = 7.4). Filamented glass capillary tubes (1.5mm/0.84mm; World Precision Instruments, USA) were pulled by a micropipette puller (Sutter, USA) prior to each experiment to pipette resistances of 3–5 MΩ for HEK 293 cells and 3–6 MΩ for cultured primary neurons. The pipettes were filled with intracellular solution (120mM K-aspartate, 4 mM NaCl, 4 mM MgCl_2_, 1 mM CaCl_2_, 10 mM EGTA, 3 mM Na_2_ATP and 5 mM HEPES, pH = 7.2) and held by a pipette holder (HEKA, Germany) mounted on a micromanipulator (Scientifica, UK). Whole cell voltage clamp and current clamp recordings of transfected cells were conducted using a patch clamp amplifier (HEKA, Germany). A holding potential of -70 mV was used for all recordings including neuronal recording with cultured neurons until switched to current clamp mode.

### Fluorescence microscopy

An inverted microscope (IX71; Olympus, Japan) equipped with a 60X oil-immersion lens with 1.35-numerical aperture (NA) was used for epifluorescence imaging. A 470nm light-emitting diode (bandwidth: 25 nm) placed in a 4-wavelength LED housing (LED4D242, Thorlabs, USA) was used for the experiments of Ace2N with Farnesylated Clover. A 4 - channel LED driver and its software (DC4100, Thorlabs, USA) were used to control the LED. For the other experiments, the light source was a 75 W Xenon arc lamp (Osram, Germany) placed in a lamp housing (Cairn, UK). CFP was imaged using a filter cube consisting of an excitation filter (FF434/17), a dichroic mirror (FF452-Di01) and an emission filter (FF01-479/40), YFP was imaged with a filter cube consisting of an excitation filter (FF497/16), a dichroic mirror (FF516-Di01) and an emission filter (FF01-535/22), CFP-YFP FRET was imaged using an optical splitter (Cairn, UK) consisting of 2 filter cubes, the first composed of excitation filter (FF434/17), a dichroic mirror (FF452-Di01) without emission filter, and the second cube consist of emission filter (FF497/16), a dichroic mirror (FF510-Di02) and emission filter (FF01-535/22). GFP-RFP FRET was imaged using 2 filter cubes, the first composed of excitation filter (FF475/23), a dichroic mirror (FF495-Di02) without emission filter, and the second cube consist of emission filter (FF520/40), a dichroic mirror (FF560-Di01) and emission filter (FF01-645/75). CFP-RFP FRET was imaged using 2 filter cubes, the first composed of excitation filter (FF434/17), a dichroic mirror (FF452-Di01) without emission filter, and the second cube consist of emission filter (FF497/16), a dichroic mirror (FF510-Di02) and emission filter (FF01-645/75) or (FF585/29) (all by Semrock, USA). Two cameras were mounted on the microscope through a dual port camera adapter (Olympus, Japan). A slow speed color charge-coupled-device (CCD) camera (Hitachi, Japan) was used to aid in locating cells and during patch clamp experiments. Fluorescence changes of the voltage indicators were typically recorded at 1 kHz frame rate by a high-speed CCD camera (RedShirtImaging, USA) unless otherwise specified. All the relevant optical devices were placed on a vibration isolation platform (Kinetic systems, USA) to avoid any vibrational noise during patch clamp fluorometry experiments.

### Data acquisition and analysis

Resulting images from patch clamp fluorometry were acquired and analyzed for initial parameters such as fluorescence change [ΔF = Fx-F0] or fractional fluorescence change values [ΔF/F =((Fx-F0)/F0) * 100] by Neuro**P**lex (RedShirtImaging, USA) and Microsoft Excel (Microsoft, USA). The acquired data from whole cell voltage clamp experiments of HEK 293 cells were averaged for 8 trials (technical replication) unless otherwise noted. The number of recorded cells in this work is biological replicates and the number of trials averaged for each HEK 293 cell should be interpreted as technical replication. Data were collected from recorded cells that did not lose their seals during the whole cell voltage clamp recording. ΔF/F values for the tested voltage pulses were plotted in OriginPro 2016 (OriginLab, USA). A ΔF/F trace versus time graph for each cell was also fitted for either double or single exponential decay functions in OriginPro 2016 as described previously (Piao et al. 2015).

## Results

### Cotransfection of InterFRET-GEVIs yields voltage-dependent optical signals

Introduction of monomer inducing mutations to the FP of ArcLight-type GEVIs resulted in a substantial reduction of the voltage-dependent optical signal (Kang & Baker, 2016). That result suggested the possibility that dimerization of the FP domain was necessary for the optical response of ArcLight. To test that hypothesis, we employed FRET imaging by replacing the SEpH FP in ArcLight with either the cyano FP (CFP), Cerulean, (Rizzo, Springer, Granada, & Piston, 2004), or the yellow FP (YFP), Venus (Nagai et al., 2002). These two constructs are denoted as ArcLight-CFP or ArcLight-YFP respectively. Figure 1 compares the optical signals from HEK cells expressing either the FRET donor construct alone, ArcLight-CFP (Figure 1A), the FRET acceptor construct alone, ArcLight-YFP (figure 1B), or the co-expression of the FRET pair (Figure 1C). Whole cell voltage clamp of HEK cells expressing only ArcLight-CFP yielded a relatively small voltage-dependent signal that reached a maximum of about 3% ΔF/F. The signal size from cells expressing only ArcLight-YFP was roughly the same but also exhibited a noticeable degree of bleaching. Cotransfection of ArcLight-CFP and ArcLight-YFP yielded a much larger optical response in which the ArcLight-YFP signal inverted its polarity. This reciprocal change in fluorescence was indicative of a FRET signal. Upon depolarization of the plasma membrane, the FRET efficiency improved as the donor chromophore fluorescence was reduced by over 10% ΔF/F/100 mV and the acceptor fluorescence was increased by 14% ΔF/F/100 mV indicating that the distance between the chromophores had decreased and/or that the orientation of the chromophores had improved during the depolarization of the plasma membrane.

**Figure 1.**
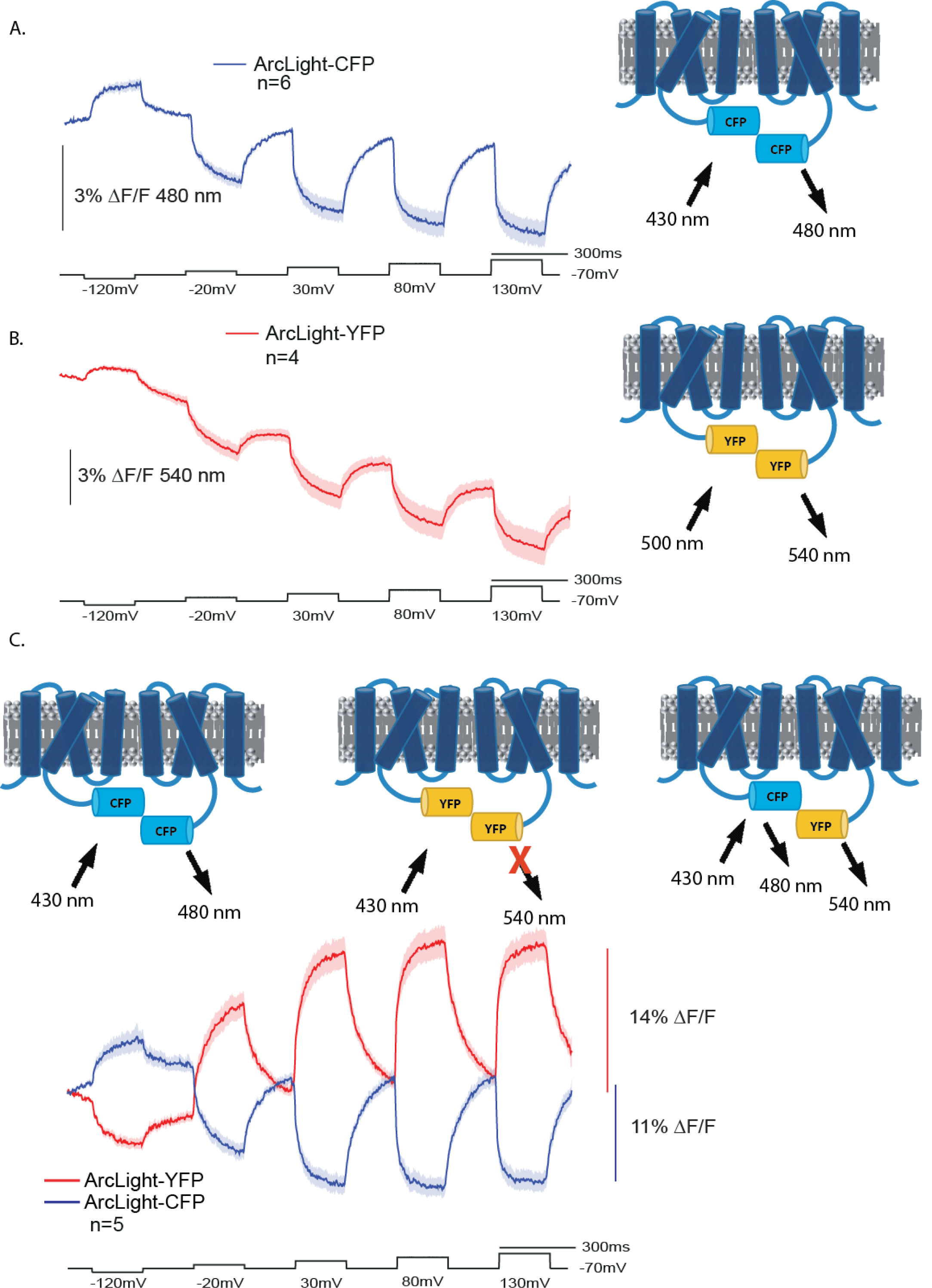
Intermolecular-FRET GEVIs. A. Optical traces from ArcLight-CFP. HEK 293 cells expressing only ArcLight-CFP were voltage clamped at a holding potential of -70 mV and subjected to a five pulse protocol (black trace). Excitation light for ArcLight-CFP was 430 nm. The optical trace in blue was acquired at 1 kHz. The solid blue line represents the mean from 6 cells. The shaded area is the standard error of the mean. B. Optical traces from ArcLight-YFP. HEK 293 cells expressing only ArcLight-YFP were voltage clamped as described in A. Excitation light was 500 nm for ArcLight-YFP. The solid red line represents the mean from 4 cells. C. Optical traces from intermolecular-FRET GEVIs. The GEVI schematics represent the three potential interactions upon co-transfection of HEK-293 cells with ArcLight-CFP and ArcLight-YFP. Excitation at 430 nm limits the contribution of the ArcLight-YFP/ArcLight-YFP interactions since the excitation peak of YFP is near 500 nm. Optical emissions at 480 nm and 540 nm were collected simultaneously with the use of an optical splitter (see methods). The blue trace is the 480 nm emission average from 5 cells voltage clamped as described in A. The red trace is the 540 nm emission average.

Since no effort was made to optimize the interaction of the molecules containing the FRET FP pairs, the optical signal in figure 1C is a mixture of ArcLight-CFP/ArcLight-YFP as well as the potential ArcLight-CFP/ArcLight-CFP and ArcLight-YFP/ArcLight-YFP associations. However, the ArcLight-CFP only signal and the ArcLight-YFP only signal are small. In addition, the wavelength of excitation in Figure 1 C was 430 nm which further reduces the ArcLight-YFP/ArcLight-YFP signal since the excitation peak for Venus is near 500 nm. Despite this imperfect system, the interFRET-GEVI signal exceeds the optical signals seen for other GEVIs that use intramolecular FRET such as Nabi (Sung et al., 2015) or VSFP-Butterfly (Akemann et al., 2012).

### The voltage sensor domain determines the speed of the optical response

The Bongwoori family of GEVIs are ArcLight-derived sensors that have faster kinetics (Lee et al., 2017; Piao, Rajakumar, Kang, Kim, & Baker, 2015). When the YFP/CFP inter-FRET version of Bongwoori-R3 is expressed in HEK 293 cells, a robust signal is again seen with faster kinetics (Figure **2**A).

**Figure 2.**
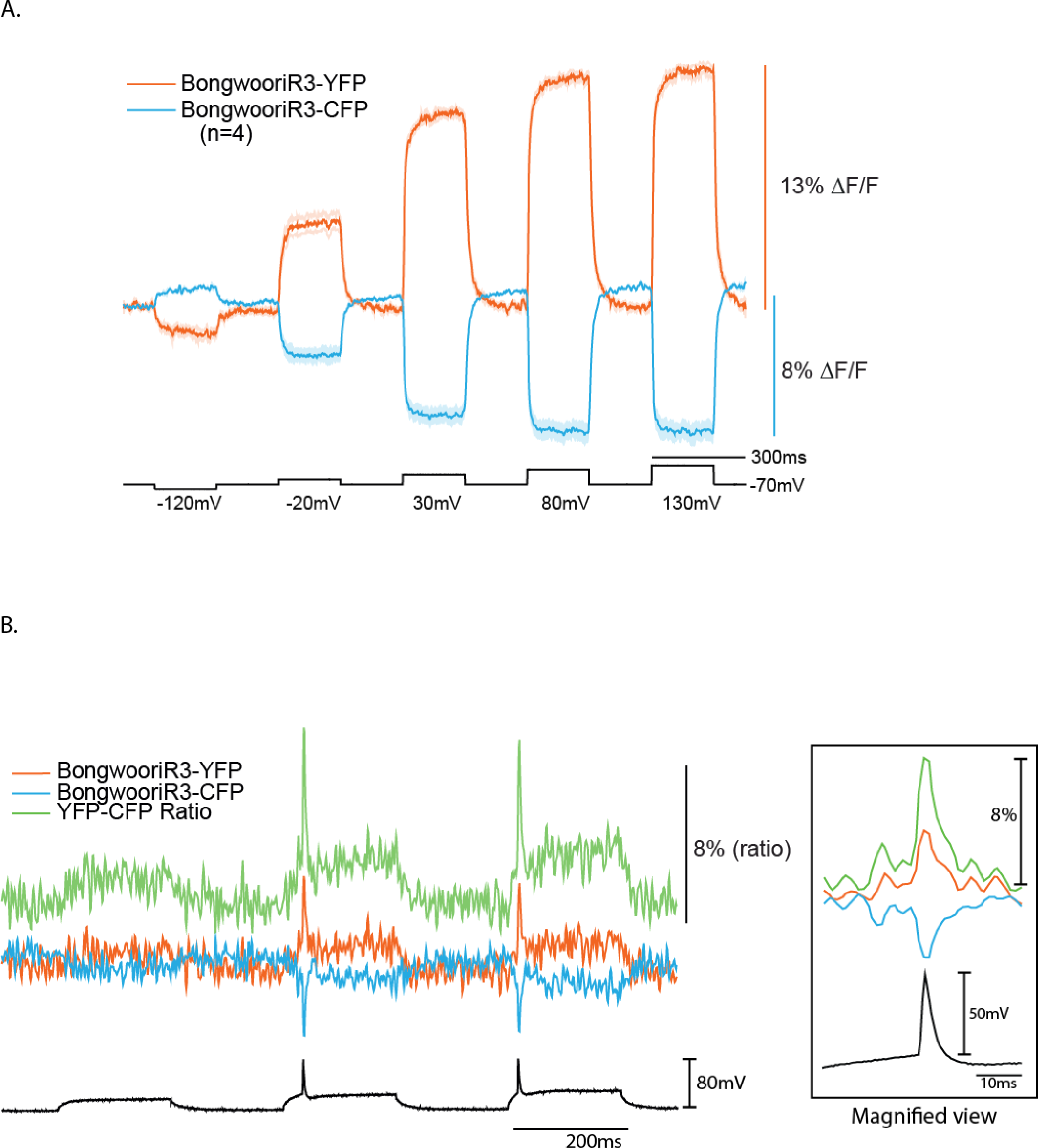
The inter-FRET signal from BongwooriR3-derived GEVIs. A. The traces are from HEK cells co-expressing BongwooriR3-Cerulean and BongwooriR3-Venus. The dark line is the mean and the shaded area is the standard error of the mean from 4 cells. The blue trace is from CFP emission at 480 nm. The red trace is from YFP emission at 540 nm. The excitation wavelength was 430 nm. B. The trace is from a hippocampal neuron in culture under whole cell current clamp co-expressing BongwooriR3-Cerulean and BongwooriR3-Venus. The black trace is voltage. The blue trace is CFP emission at 480 nm. The red trace is from YFP emission at 540 nm. The green trace is the ratio of YFP/CFP fluorescence. Inset is a time expanded view of the first action potential.

Since this version is faster, we also expressed this FRET pair in cultured hippocampal neurons resulting in a fast, ratio-metric signal during action potentials (Figure **2**B).

### Monomeric mutations to the FP do not abolish the voltage-dependent FRET signal

Most derivatives of GFP from *A. Victoria* exist as protein dimers. An exception is CFP due to the N146I mutation rotating the 7th β-strand and exposing the side chain of Y145 (numbering based on GFP amino acid sequence) which interferes with dimer formation (von Stetten, Noirclerc-Savoye, Goedhart, Gadella, & Royant, 2012). However, since CFP can form dimers in the mM range (Espagne et al., 2011), the effective protein concentration at the plasma membrane may be sufficient for some dimerization since diffusion is limited to 2-dimensional space. We therefore introduced monomeric favouring mutations to the FPs of the ArcLight-FRET pairs.

The three mutations to GFP that favour the monomeric form are A206K, L221K, and F223R (Zacharias, Violin, Newton, & Tsien, 2002). The L221K mutation exhibited very little effect on the voltage-dependent FRET signal (Figure 3). The YFP signal was virtually the same (14% compared to 13%), and the CFP signal was slightly diminished (8% compared to 11%). The A206K mutation which is the most common monomeric mutation slightly affected the YFP signal (10% compared to 14%) but nearly cut the CFP signal in half (6% compared to 11%). The F223R mutation had the largest effect reducing the YFP signal from 14% to below 4% and the CFP signal from 11% to 2%.

**Figure 3.**
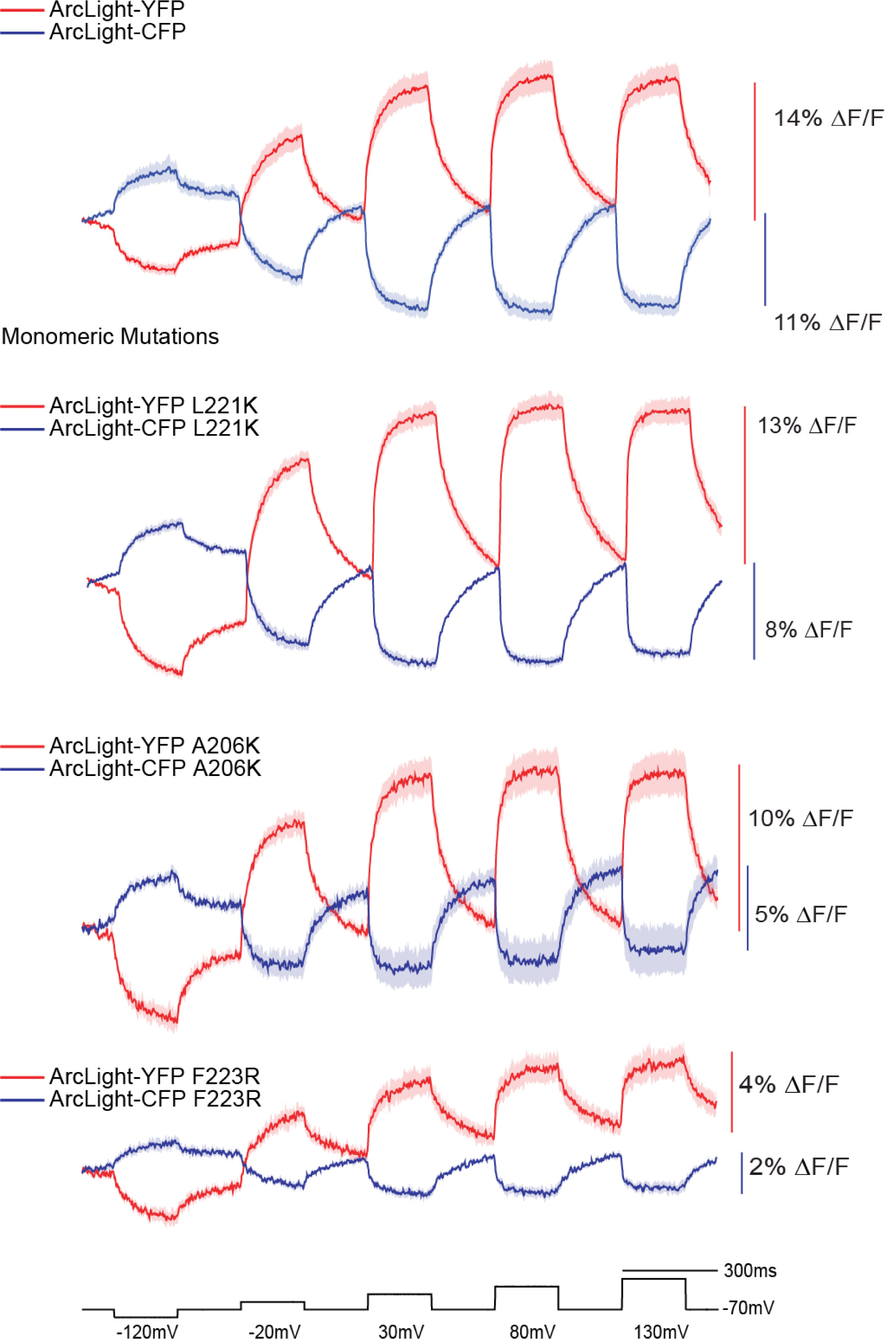
Monomeric mutations to the FP domains may reduce but do not destroy the voltage-dependent, intermolecular-FRET signal. The top trace is the intermolecular-FRET signal reproduced from figure 1C for comparison purposes. Bottom three traces are from HEK cells expressing ArcLight intermolecular FRET pairs containing one of the three FP monomeric mutations (A206K, L221K, or F223R). The dark trace is the mean from at least 4 cells. The shaded area is the standard error of the mean.

This large variation in the effects on the voltage-dependent optical signal suggests there may be some dimerization of the FP domain at the plasma membrane. However, the monomeric mutations were unable to completely destroy the optical signal suggesting that other FPs from different organisms with different wavelengths could also be used for interFRET-GEVI development.

### Expanding the spectrum and the Stokes shift of the interFRET-GEVI signals

The ArcLight family of GEVIs utilize the voltage sensing domain from the gene family of voltage-sensing phosphatases. Recently, it has been shown that the voltage sensing phosphatase can dimerize (Rayaprolu, Royal, Stengel, Sandoz, & Kohout, 2018). Dimerization via the voltage sensing domain could explain why monomeric mutations to the FP domain failed to eliminate the voltage dependent optical signal. If the voltage sensing domain was responsible for dimerization, that would enable the use of any FRET pair for imaging voltage.

Replacement of the SEpH FP with the red-shifted FP, mRuby2 (Lam et al., 2012), again yielded a GEVI with a modest voltage dependent signal of under 3% ΔF/F/100 mV when expressed in HEK 293 cells (Figure 4A). To facilitate the FRET signal for the FP, mRuby2, we also replaced SEpH in ArcLight with the GFP-derived FP, Clover (Lam et al., 2012). HEK 293 cells expressing only ArcLight-Clover yielded an even smaller voltage-dependent signal below 2% ΔF/F/100 mV (Figure 4B). However, when ArcLight-Clover and ArcLight-mRuby2 were cotransfected into HEK 293 cells, a much improved voltage-dependent signal was observed (Figure 4C). The ArcLight-Clover signal decreased by 7% during a 100 mV depolarization step while the ArcLight-mRuby2 signal increased by nearly 11%. Like the CFP/YFP version, depolarization of the plasma membrane increased the FRET efficiency indicating that the chromophores of the FPs were getting closer and/or moving into a better orientation for energy transfer.

**Figure 4.**
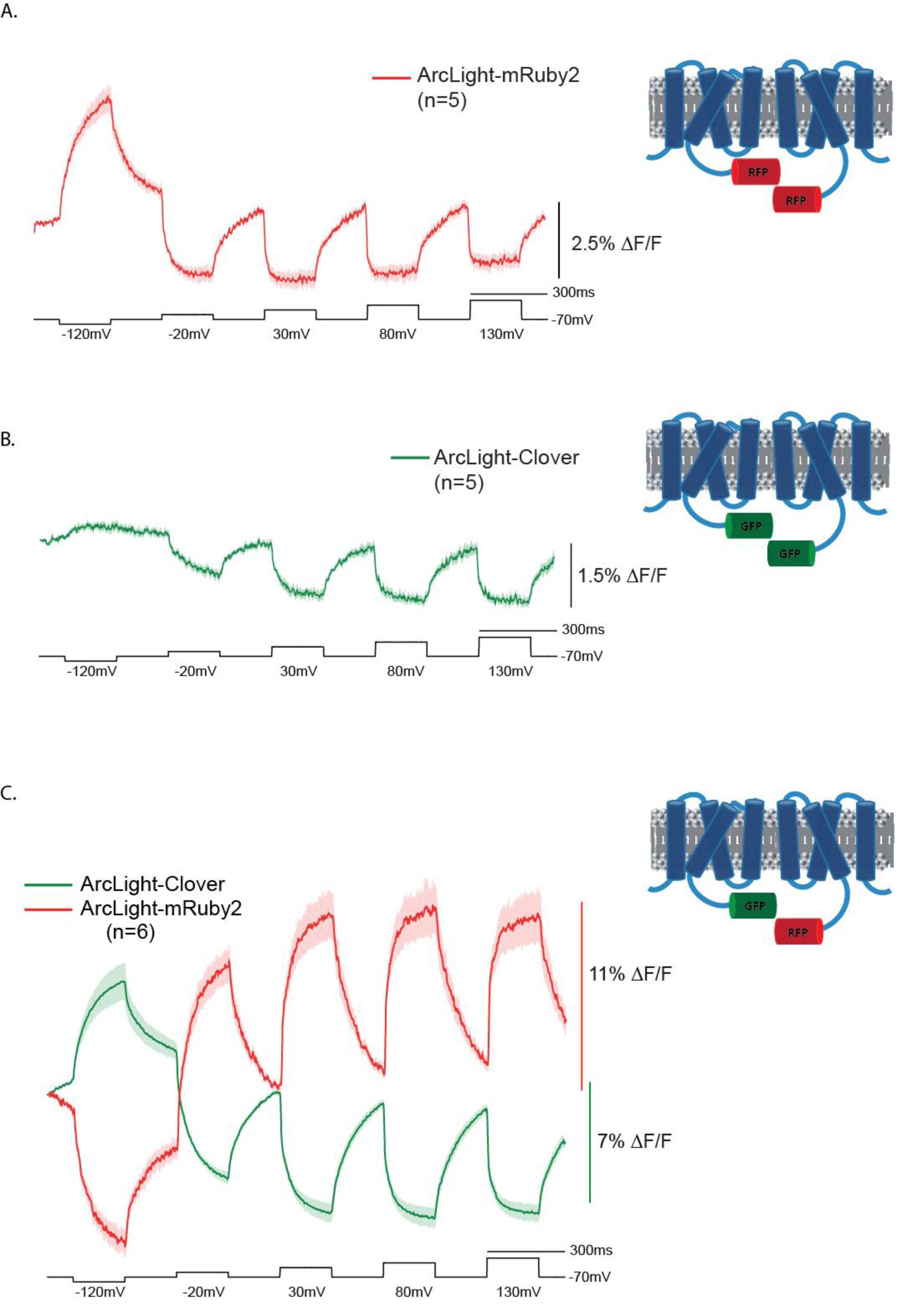
Green/Red Inter-FRET GEVI voltage dependent signals. A. The trace is the average of 5 HEK cells expressing ArcLight-mRuby2. B. The trace is the average from 5 HEK cells expressing ArcLight-Clover. C. The traces are from 6 HEK cells co-expressing ArcLight-Clover/ArcLight mRuby2. The voltage steps are in black. In all cases the dark trace is the mean while the shaded area is the standard deviation of the mean.

### Investigating the multimeric nature of ArcLight-derived GEVIs enabled a 200 nm Stökes shift

The rather robust FRET signals for both the CFP/YFP pair and the GFP/RFP pair indicated that the FP domain of ArcLight-derived GEVIs are in close proximity which suggested that the association was occurring at least in part via the voltage sensing domain. To determine the extent of a possible multimerization, we attempted an inter-FRET from CFP to RFP via YFP. Triple transfection of HEK 293 cells with the ArcLight GEVI fused to CFP, YFP, and RFP resulted in a voltage-dependent signal that exhibited a Stökes shift of 215 nm (Figure 5A, left). A similar optical signal was observed when the RFP, mRuby2, was replaced with dTomato (Figure 5A, right). Using the VSD of BongwooriR3 also showed a large Stokes shift with improved speed and a more positive voltage shifted response (Figure 5A, right and 5B).

**Figure 5.**
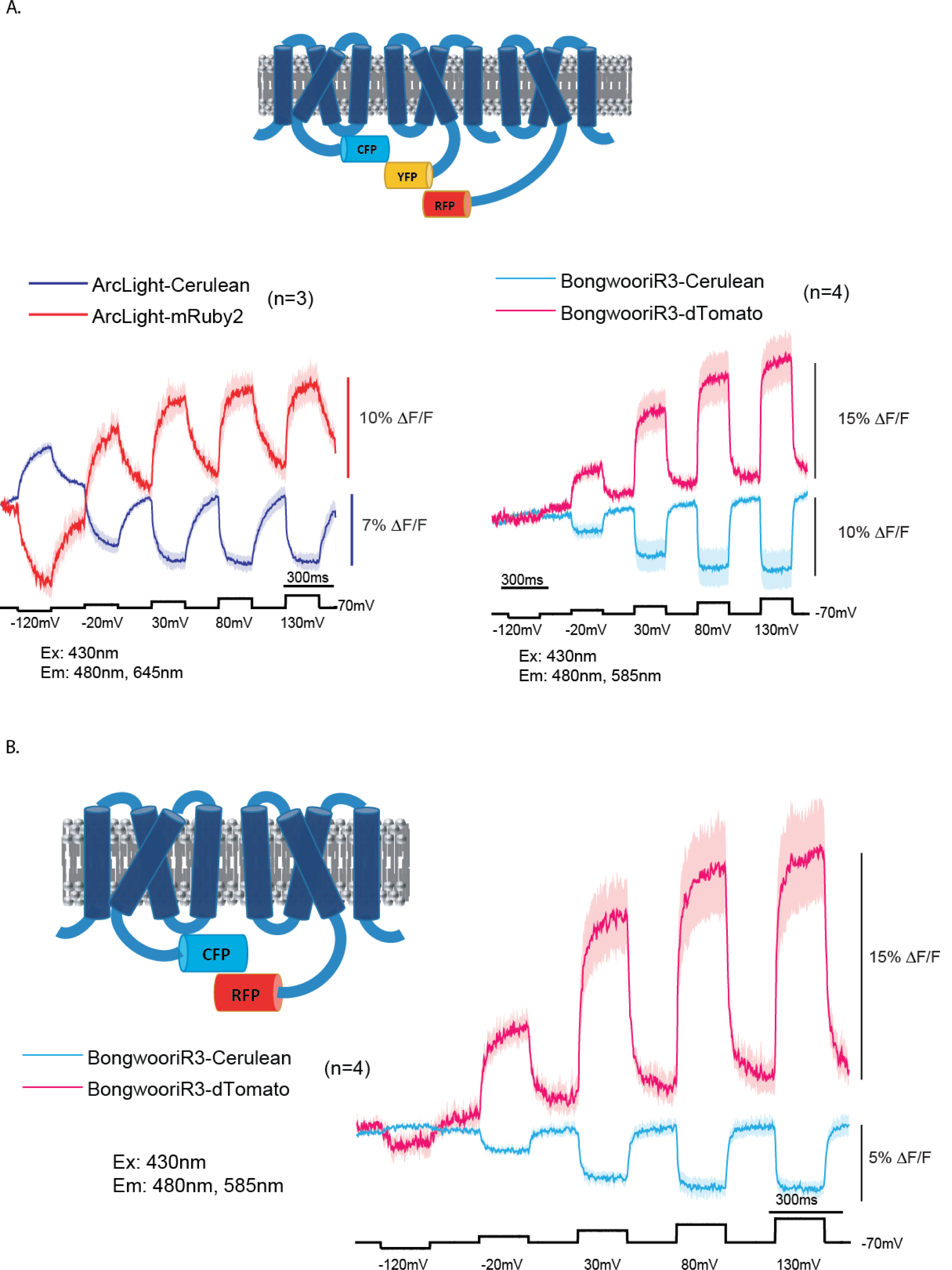
The proximity of adjacent chromophores enables a large Stökes shift. A. Voltage dependent optical responses of cells transfected with three GEVIs. The traces on the left are from HEK cells expressing ArcLight-Cerulean, ArcLight-Venus, and ArcLight-mRuby2. The blue trace is from the CFP emission at 480 nm. The red trace is from the RFP emission at 645 nm. Excitation wavelength was 430 nm. YFP emission was not recorded as we can only simultaneously record two wavelengths. The traces on the right are from HEK cells expressing the trio of GEVIs, Bongwoori-Cerulean, Bongwoori-Venus, and Bongwoori-dTomato. The blue trace is from the emission of CFP at 480 nm. The red trace is from the emission of RFP at 585 nm. Excitation wavelength was also 430 nm. B. Voltage dependent optical signals of HEK cells transfected with BongwooriR3-Cerulean and BonwooriR3-dTomato. The blue trace is from the CFP emiision at 480 nm. The red trace is from the RFP emission at 585 nm.

To determine if the large Stökes shift was due to a triple FRET signal, we also imaged HEK cells expressing only the inter-FRET CFP/RFP Bongwoori-R3 constructs (Figure 5B). Surprisingly, excitation at 420 nm yielded a voltage-dependent optical signal at 585 nm without the presence of YFP. Despite the poor overlap of Cerulean’s emission profile to dTomato’s excitation spectra, Cerulean has a broad emission shoulder enabling FRET from Cerulean to dTomato. Comparison of the signals from HEK cells expressing the triple FRET constructs (CFP/YFP/RFP) to HEK cells only expressing the CFP/RFP versions showed a similar dynamic range for RFP (20% ΔF/F/200 mV), but the CFP signal for the triple FRET expressing cells was twice that of cells not expressing the YFP version. This disparity could be the result of CFP directly transferring energy to RFP in both experiments. Assuming that only dimers formed in cells expressing all three FP constructs (CFP/YFP/RFP), the CFP signal would be a culmination of the voltage dependent quenching from the CFP/CFP interaction, the CFP/YFP association as well as the CFP/RFP interaction. In the CFP/RFP experiment, the CFP signal would consist of only CFP/CFP and CFP/RFP interactions.

### Monitoring the movement of a single FP using heterogeneous VSDs

The movement of S4 has been shown to mediate the voltage dependent optical signal for ArcLight-derived GEVIs (Kang & Baker, 2016; Treger, Priest, & Bezanilla, 2015; Villalba-Galea et al., 2009). One potential reason the intermolecular FRET signal is relatively larger than earlier FRET constructs (Akemann et al., 2012; Dimitrov et al., 2007; Perron et al., 2009; Sung et al., 2015) is that the voltage-induced conformational change is amplified. Since both the donor and acceptor FPs are moving, the net distance/orientation change may be greater. Previous FRET GEVIs incorporated both the donor and acceptor FPs in the same molecule thereby limiting the change in distance/orientation.

To visualize the individual contribution of the FRET donor or FRET acceptor on the voltage signal, we utilized combinations of voltage sensing domains with varying sensitivities. An inter-FRET GEVI pair that responds to very positive membrane potentials will not yield an optical signal during a small depolarization step (figure 6A). The GEVI, CC1, is an ArcLight-derived GEVI that has a positively shifted voltage range (Piao et al., 2015). The V_1/2_ (the membrane potential that corresponds to one half of the total fluorescence change) for CC1 is nearly +100 mV. An inter-FRET GEVI pair that responds to more negative potentials will yield an optical signal for smaller depolarization steps (Figure 6B). The V_1/2_ for ArcLight is around -20 mV. By combining these different voltage sensing domains, we could observe the motion of one or both S4 segments depending on the size of the depolarization step (Figure 6C).

**Figure 6.**
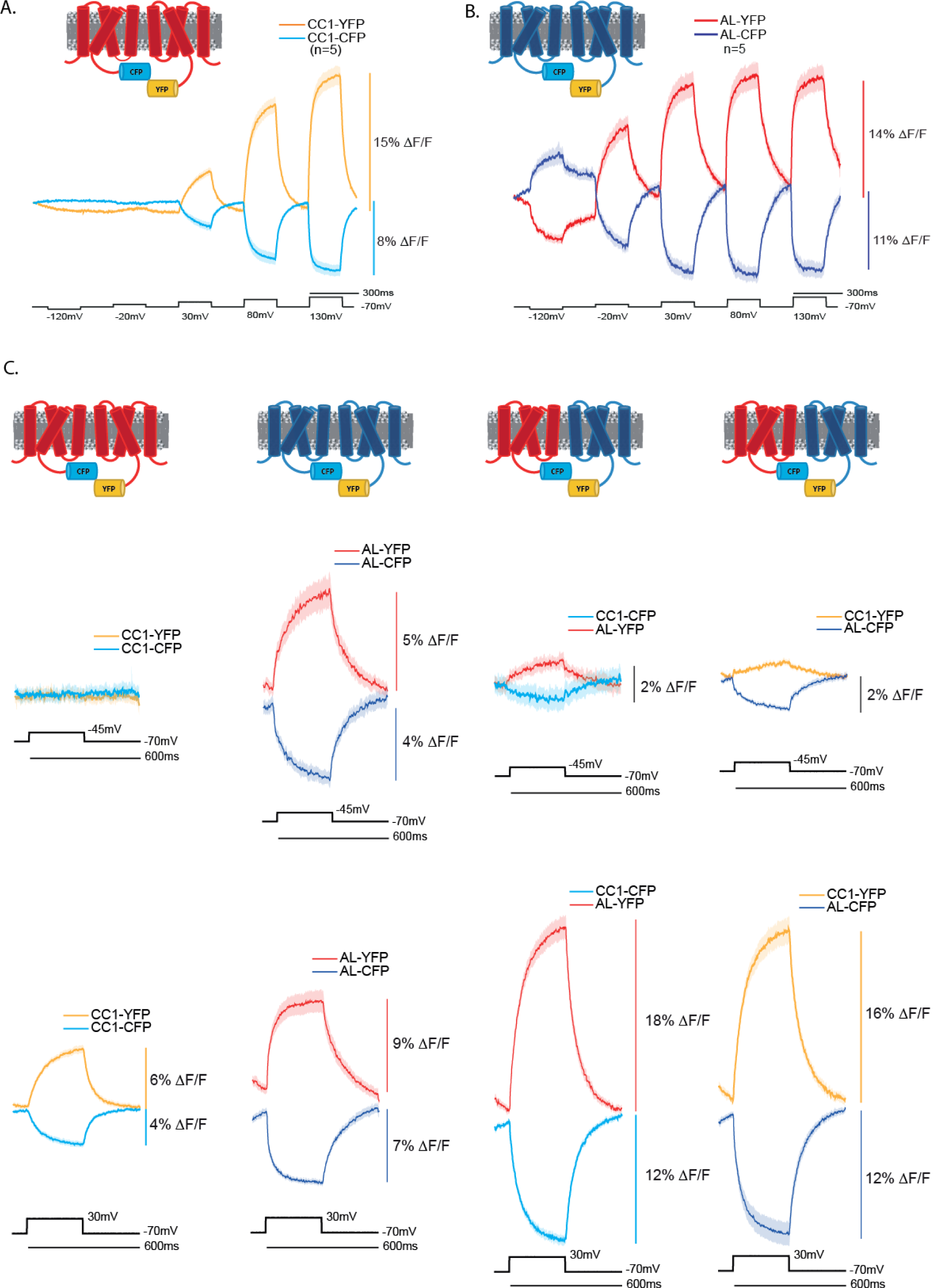
Decoupling the voltage dependent FRET signal by varying the voltage sensitivities of the voltage sensing domains. A. An inter-FRET GEVI pair using the voltage sensing domain from CC1 (red transmembrane segments) which has positively shifted voltage responsive range. The donor fluorescence trace is in light blue (CC1-CFP). The acceptor fluorescence trace is in yellow (CC1-YFP). Voltage steps are in black. B. An inter-molecular FRET GEVI using the voltage sensing domain from ArcLight (blue transmembrane segments) which responds to more negative membrane potentials. The donor fluorescence trace is in dark blue (AL-CFP). The acceptor fluorescence trace is in red (AL-YFP). C. Four columns of the different combinations of CC1 and ArcLight inter-FRET GEVI pairs. The top row of traces are from HEK cells experiencing a 25 mV depolarization of the plasma membrane potential. The bottom row of traces are from HEK cells experiencing a 100 mV depolarization of the plasma membrane potential. The colors of the traces and voltage sensing domains are as described in A. and B. Excitation wavelength was 430 nm for all experiments. CFP emission was measured at 480 nm and YFP emission was measured at 540 nm.

When the donor and acceptor FPs are fused to the CC1 voltage sensing domain, there is no observable signal when the membrane potential of HEK cells is depolarized by 25 mV from a holding potential of -70 mV. In contrast, when the inter-FRET GEVI pair both utilize the voltage-sensing domain from ArcLight, a slow but obvious FRET signal is detected during a 25 mV depolarization (Figure 6C). When the membrane potential is depolarized by 100 mV to +30 mV, both the CC1 FRET pair and the ArcLight FRET pair yield signals (Figure 6C). Since only the voltage sensing domain from ArcLight responds to the 25 mV voltage step, we then tested inter-FRET GEVIs consisting of CC1-CFP and ArcLight-YFP as well as ArcLight-CFP and CC1-YFP. When the positively shifted CC1 voltage-sensing domain is fused to the FRET donor and the FRET acceptor is fused to the ArcLight voltage sensing domain, a small signal around 1% ΔF/F was observed during the 25 mV depolarization step. This was less than when both FRET FPs are attached to the ArcLight voltage-sensing domain since only one of the two S4 segments was responding.

However, when the plasma membrane is depolarized by 100 mV, the signal is nearly twice the size as seen when both FRET pairs are fused to either the CC1 or the ArcLight voltage sensing domain. This result may be due to the difference in the number of amino acids in the linker segment between the voltage sensing domain and the FP. For ArcLight, there are 11 amino acids in the linker segment while CC1 has 22 amino acids in the linker segment. The movement of the FP may also be different for the two constructs which might improve the FRET efficiency for the CC1/ArcLight pairing during stronger depolarization of the plasma membrane.

### Monitoring the movement of a single FP using lipid anchors

A recurring problem when monitoring inter-FRET GEVI signals is the contamination of the signal caused by homodimers of the FRET donors. While these signals are small (Figures 1A, 1B, and 4), they can still be seen in some recordings. For instance, in figure 6C when the donor FP is fused to the ArcLight and the acceptor FP is fused to the positively shifted CC1 voltage sensing domain, the on and off rates for the donor FP do not mirror the rates for the acceptor FP. This is because the donor FP signal is contaminated with the ArcLight-CFP/ArcLight-CFP voltage dependent signal. This is not the case when the donor FP is fused to positively shifted CC1 construct. There still exists CC1-CFP/CC1-CFP associations but since CC1 does not respond to that voltage, the FRET signal is less contaminated resulting in a reduction of the ΔF/F in the CFP channel for the CC1-CFP/ArcLight-YFP pairing. There may still be a contaminating signal from ArcLight-YFP/ArcLight-YFP even though the excitation wavelength was 430 nm.

To remove the contaminating signal from homodimers, we replaced one of the FRET FPs with a farnesylated version lacking a voltage sensing domain (Figure 7). When the FRET donor FP, CFP, is fused to the Bongwoori-R3 VSD and co-expressed in HEK cells with a farnesylated YFP (Figure 7A), a 2% ΔF/F for YFP can be seen when the plasma membrane is depolarized by 200 mV. The 200 mV depolarization step results is a 3% ΔF/F signal in the CFP channel. When the FRET pair is reversed by fusing the donor FP to the VSD of Bongwoori-R3, there is still a voltage-dependent FRET signal (Figure 7B).

**Figure 7.**
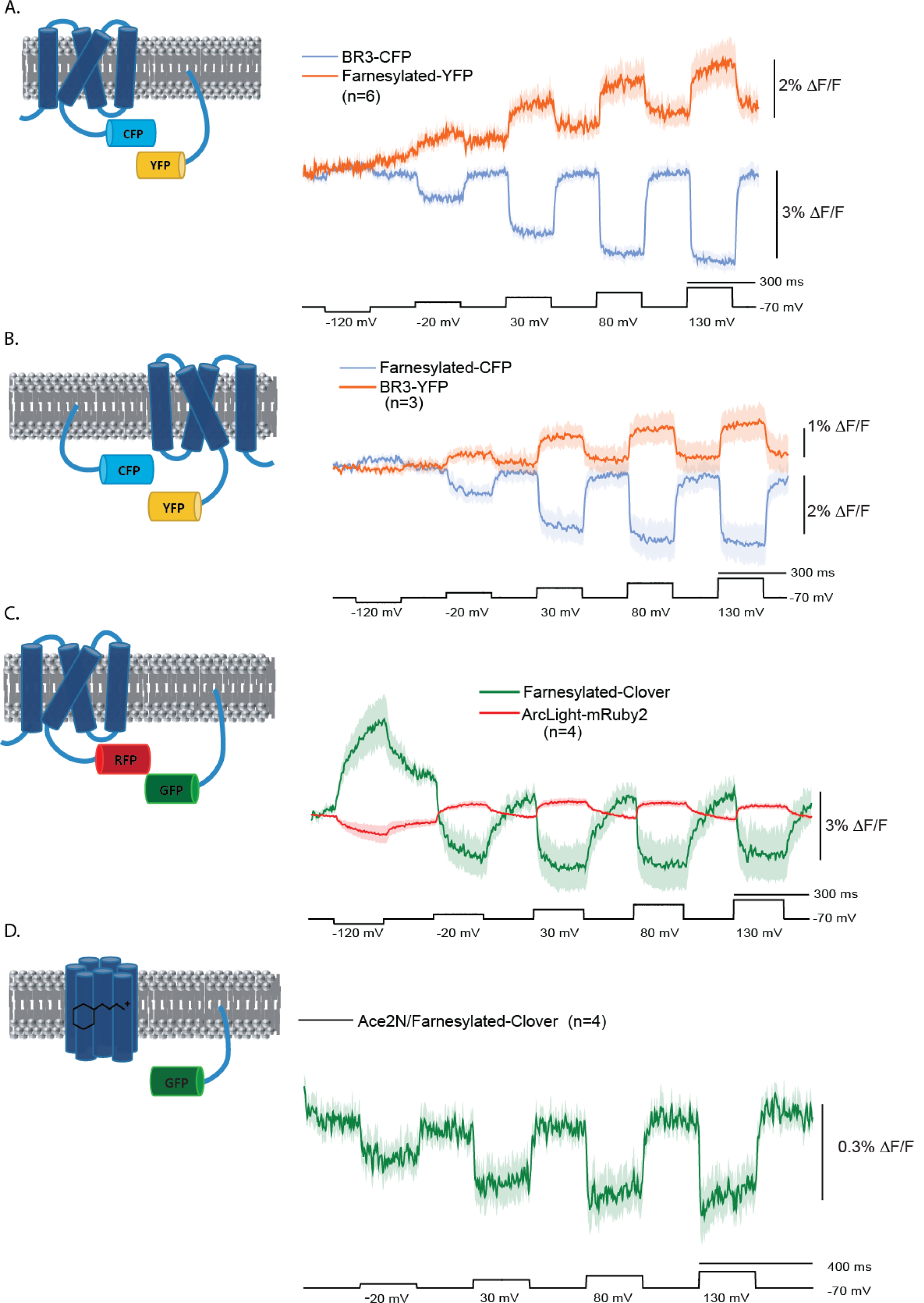
Intermolecular FRET with a farnesylated FP. A. CFP fused to the voltage sensing domain of the GEVI, Bongwoori-R3, was co-expressed in HEK cells with a farnesylated version of YFP. The blue trace is the emission at 480 nm. The red trace is the emission at 540 nm. The excitation wavelength was 430 nm. Dark line is the mean of 6 cells. Shaded areas are the standard error of the mean. B. YFP fused to the voltage sensing domain of Bongwoori-R3 was co-expressed with a farnesylated CFP. The color coding and wavelengths are as in A. C. The FP, mRuby2, was fused to the GEVI, ArcLight and co-expressed with a farnesylated Clover FP. The green trace is the emission at 520 nm. The red trace is the emission at 585 nm. The excitation wavelength was 470 nm. D. Intermolecular FRET of a rhodopsin GEVI. Ace_2N (Gong et al., 2015), lacking the mNeon FP was co-expressed with farnesylated-Clover.

A farnesylated GFP, Clover, was also able to yield a voltage dependent FRET signal when co-expressed with the GEVI ArcLight-mRuby2 (Figure 7C). Again this signal is much reduced than when both FP are fused to different VSDs which may indicate that the VSD enhances the association of intermolecular FRET partners. However, we have not tried to optimize the linker link between the FP and the VSD for all of these associations which may also contribute to the dynamic fluorescent signal.

Intermolecular FRET also works for the rhodopsin family of GEVIs (Figure 7D). When HEK cells were co-expressed with a farnesylated Clover FP and the rhodopsin GEVI, Ace2N taken from Ace2N_mNeon (Gong et al., 2015) lacking the mNeon green domain, a voltage-dependent optical signal was observed. While this signal is significantly reduced, it provides a proof-of-principle that intermolecular FRET can potentially improve the flexibility for rhodopsin-based GEVIs as well.

## Discussion

The experimental paradigm of intermolecular FRET using ArcLight-derived GEVI**s** has revealed new information about the dimerization mechanism coupling voltage to fluorescence. The suggested homodimer fluorescence mechanism of ArcLight-derived GEVIs that interact via their cytoplasmic FP domains was expanded by the discovery of voltage dependent intermolecular FRET signals (Figures 1 and 2). The observation that monomeric mutations to the FP dramatically reduced the voltage dependent optical signal of ArcLight suggested that the association was primarily via the FP domain (Kang & Baker, 2016). However, the intermolecular FRET signals have revealed that the interaction between the FPs of adjacent GEVIs persists despite the presence of monomeric mutations to the FP domain (Figure 3) suggesting other regions of interaction. A recent study reported that the intact voltage sensing *Ciona intestinalis* phosphatase, Ci-VSP, can dimerize (Rayaprolu et al. 2018). Since the voltage sensing phosphatase lacks an FP domain and the intermolecular FRET GEVIs lack the phosphatase domain, the voltage sensitive domain is likely responsible at least in part for dimerization.

The association of the voltage sensitive domain enabled the discovery of a new family of intermolecular FRET GEVIs by fusing the donor FP to one VSD and the acceptor FP onto a separate VSD. We can now create GEVIs that 1) get brighter in response to depolarization, 2) have inverted signals that can be used to remove motion artefacts, 3) improve the dynamic range by amplifying the voltage-induced conformational change, 4) exploit multiple wavelengths, greatly increasing the Stökes Shift, 5) permit the development of better 2-photon voltage sensors, 6) restrict expression to more precisely defined cellular populations.

ArcLight has a large dynamic response to membrane potential changes and gets dimmer upon depolarization of the plasma membrane (Jin et al., 2012). In an independent comparison of GEVIs from 2019, ArcLight was the only GEVI capable of reporting neuronal activity in response to visual stimuli in the mouse visual cortex (Bando et al., 2019). Unfortunately, the signal to noise ratio is low for several reasons. One is that the optical responsive protein is limited to the plasma membrane which generates a high background fluorescence since dendrites as well as axons are fluorescent (Nakajima & Baker, 2018). Another is that the voltage change experienced by a neuron can vary from small depolarizing or hyperpolarizing synaptic potentials that last tens of milliseconds to large voltage changes from action potentials that lasts one to two milliseconds. One way to reduce the high background is to create a GEVI that starts dim and gets brighter upon depolarization. Several rhodopsin-based GEVIs exhibit this characteristic but require intense illumination which limits the number of cells that can be imaged simultaneously (Adam et al., 2019; Lou et al., 2016). The rhodopsin-based GEVIs also respond poorly upon 2-photon illumination and exhibit light induced currents as well as chromogenic effects. Two ArcLight-derived GEVIs have been developed that also become brighter upon depolarization of the plasma membrane, FlikR1 which has the added advantage of being red-shifted (Abdelfattah et al., 2016) and Marina (Platisa, Vasan, Yang, & Pieribone, 2017). Both show promise but have not yet exhibited the large change in fluorescence that ArcLight has.

Intermolecular FRET GEVIs offer a new solution to this problem. Depending on the emission wavelength being monitored, the voltage-induced optical signal can start bright and get dimmer during depolarization, or start dim and get brighter (Figures 2 and 4). With the use of an optical splitter, both emission wavelengths can be monitored simultaneously enabling the removal of motion artefacts. Motion artefacts will affect the fluorescence in both emission wavelengths similarly while a voltage-induced change would generate opposite fluorescent signals in the two emission channels. In addition, the inter-FRET GEVIs can utilize any FRET pair, CFP/YFP (Figures 1,2, and 3), GFP/RFP (Figure 4), and even the poor FRET pair of CFP/RFP (Figure 5). In the case of the CFP/RFP pairing, the Stökes Shift was over 200 nm which we believe is a record for GEVIs. While this may not be an ideal pairing (the transfer efficiency may be too low), having the emission wavelength get brighter upon depolarization and be 200 nm away from the excitation wavelength would be an ideal situation for in vivo recordings. In the future, we plan to expand the repertoire of FPs to include those that are very bright under 2-photon illumination for use as the inter-FRET GEVI donor in combination with an appropriate FP acceptor.

Another advantage of inter-FRET GEVIs is that the dynamic range is improved since the conformational change is the result of the movement of two VSDs (Figure 6). Having the donor and acceptor FPs on separate proteins enables the mixing of heterogeneous voltage sensitive domains. By combining a voltage sensing domain that only responds to very positive potentials (Figure 6A) with a voltage sensing domain that responds to physiologically relevant voltages (Figure 6C), an experimenter can control the movement of one or both by clamping the plasma membrane to different potentials. To verify that the movement of only the FRET donor or only the FRET acceptor could elicit a voltage-dependent optical signal, co-expression of inter-FRET GEVIs with a farnesylated partner lacking a voltage sensing domain were also able to respond to potential changes (Figure 7). The signal size of inter-FRET GEVIs partnered with a farnesylated FP were reduced roughly five fold suggesting that two VSDs improves the association of the GEVIs and/or that the linker length for the farnesylated-FP pairing is not optimal. Despite the reduced optical signal, the fact that a voltage-dependent FRET signal can occur between two membrane proteins creates the possibility that any membrane protein could be used as a potential FRET partner. The voltage-dependent FRET signal could then be limited to different parts of the cell like the axon or synapse. This may also be true for the electrochromic FRET GEVIs that utilize bacterial rhodopsin such as Ace2N-mNeon (Gong et al., 2015).

Inter-FRET GEVIs also have the potential advantage of finely tuning the neuronal cell type expressing both the donor and acceptor FRET GEVIs. By placing the expression of the FRET donor GEVI under one promoter and the expression of the FRET acceptor GEVI under another promoter, only cells capable of expressing both would yield a voltage dependent FRET signal. While this could revolutionize the decoding of neuronal circuits, a big hurdle preventing the current use of inter-FRET GEVIs *in vivo* is the need to proportionally express both GEVIs at levels facilitating heterodimer formation. In culture, this is relatively easy as both constructs are expressed at high levels. Indeed, we did not need to optimize transfection levels to observe inter-FRET GEVI signals. However, i*n vivo*, this is not a trivial issue. This problem with variable expression is compounded with the formation of homodimers. Once the dimer site is identified, homodimers can be disrupted. Further modifications of the dimer may facilitate only heterodimer formation. Once these challenges are met, the advantages of inter-molecular FRET GEVIs should be realized.

## Acknowledgments

We thank Lawrence Cohen and Yunsook Choi for critical review of the manuscript. This report was funded by the Korea Institute of Science and Technology grants 2E26190, 2E26170, 2E30070

